# Continuous quasi-attractors dissolve with too much – or too little – variability

**DOI:** 10.1101/2023.08.16.553619

**Authors:** Francesca Schönsberg, Rémi Monasson, Alessandro Treves

## Abstract

Hippocampal place cells in bats flying in a 200m tunnel have been shown to be active at multiple locations, with considerable variability in place field size and peak rate. We ask whether such disorderly representation of one’s own position in a large environment could be stored in memory through Hebbian plasticity, and be later retrieved from a partial cue. Simulating an autoassociative network in which similarly variable place fields are encoded with a covariance rule, we find that it may serve spatial memory only within a certain variability range, in particular of field width. The working range is flanked by two dysfunctional regions, accessed through apparent phase transitions. For a large network, phase boundaries can be estimated analytically to depend only on the number of fields per cell in one case, and to be a pure number in the other, implying a maximal size of the environment that can be stored in memory.

## 1. Introduction

Much of our intuition about spatial representations in the hippocampus has been shaped by recordings of place cells in simple laboratory environments of limited extent. It has been difficult, in particular, to dispel the notion that a place cell is active, typically, in a single place field, even though it has long been observed that, especially in larger environments or in the dentate gyrus, multiple fields are common [1, 2, 3, 4, 5, 6]. Even more resilient has been the concept of a characteristic size of the place field, notwithstanding the clarification that this would in any case depend on the position of the place cell along the longitudinal extent of the hippocampus [7, 8, 9]. Generations of associative memory network models, for over a quarter of a century [10, 11, 12], have assessed the possibility that a representation of space expressed by the place cells may be encoded and later retrieved, predicated on cells expressing (usually single, if any) fields of *standard* size.

Recent studies have quantified the distribution, along relatively long trajectories, of the fields expressed by individual CA1 place cells, in rats [13], in mice [14] and in bats [15]. They include several examples of cells with over 10 fields, placed at seemingly irregular locations along a 48m track in the case of rats, or a 200m flight tunnel in that of bats. Particularly the study in bats points also at the huge variability in the peak firing rate and in the width of these multiple fields. The latter is reported to be well fit by a log normal distribution, which allows for small fields from under a meter wide, up to large ones of tens of meters. A multi-field multi-scale *neural code* is argued to be advantageous in terms of neurons required and of decoding error [15]; but can such a disorderly arrangement of place fields be the basis for a stable *memory* representation of the environment?

Theoretically, the assumption of standard fields facilitates conceptualizing a recurrent neural network as having encoded, given Hebbian learning on its synaptic weights, a continuous attractor, such that each point on the manifold is a fixed point of the dynamics and corresponds to a point in real space, whether in one, two or three dimensions (or equivalently in abstract continuous spaces of any dimensionality). Of course the fixed points would be only marginally stable, susceptible to even minimal forces acting along the manifold, and then true continuity would only arise in remote mathematical limit conditions. A real system would only have a discrete number of genuine fixed points, a bit like the bed of a river, neither exactly horizontal nor completely smooth, would begin to present small fragmentary puddles when the water level is dangerously low – but in normal conditions for all practical purposes it could serve to effectively direct the flow of water.

Much work, since the introduction of such continuous attractor neural networks (cANN) to neuroscience by [16], has been devoted to discuss what can contribute to drive their neural dynamics, also considering that they are endowed, in practice, with only a discrete number of true fixed points, rather than a really continuous attractive manifold. “Bumps” of neural activity may slide along an embedding manifold towards these fixed points due to *fast neural noise* [16, 17, 18, 19, 20, 21, 22, 23, 24, 25, 26], or driven by *firing rate adaptation* [27, 28] or by *quenched disorder* induced either by the storage of multiple maps [29, 30, 31, 32], or just by random noise [19, 33, 34, 35, 36, 37, 38]; by the *encoding of additional variables* [39, 40, 41, 42] or of a consistent *direction of motion*, in the synaptic weights [19, 43]. All these considerations may be regarded as stretching a bit the notion of continuous attractors, but still the cANN ideal provides us with a valid reference point.

Here, however, we find that variability in field size and peak rates can tear that reference apart, and reduce it to shreds; moreover, this is predicted to occur very close to the regime in which a real network is experimentally observed to operate; further, given multiple fields per unit, the ideal cANN notion is destroyed also by too little variability; finally, the shrinking domain of validity of the cANN idealization is well described by surprisingly simple analytical formulae of somewhat unclear origin.

We define a model based on the study by [15], assuming that the place field statistics they observe in hippocampal area CA1 would be similar, modulo parameter differences (e.g., sparser fields [44, 45]), in CA3. As a prevailingly recurrent network, CA3 is taken to be the main locus of David Marr’s *collateral effect* promoting memory retrieval [46, 47]. Further, we assume that the place field statistics observed during bat flight are representative of the representations to be deposited in memory, not of those that can be later retrieved in the near absence of sensory inputs. Finally, as a non-essential constraint to be relaxed later, we assume direction-invariant place fields along a 1D track or flight tunnel, which present computational advantages in a network model, including being compatible with symmetric models of synaptic plasticity, such as the covariance rule [48], and enabling a free-energy analysis [49]. Experimentally, recorded fields are directional, i.e., different in the two directions of motion, but they tend to become direction-invariant in higher dimension [50, 51] (perhaps by fusing direction-dependent fields through associative plasticity, as envisioned by [52]).

## 2. Results

### 2.1 Model definition

We first consider a mathematical network model fairly close to the statistics observed in bats by [15], with two main ingredients which are obviously not in their data:

- recurrent connections among the units (known to be abundant in CA3, not in CA1).
- Hebbian plasticity on those connections, through which the network is assumed to have stored a representation of a tunnel of length *L* – and only of that tunnel.

Pyramidal cells are modelled as threshold-linear units [53], which enable both plausible statistics and mathematical analysis.

We denote by *η*_*i*_(*s*) the recorded activity of cell *i* when the bat is at position *s* in the tunnel. This space-dependent activity is assumed to give rise, through a learning phase not explicitly modeled here, to the connections between the cells through a Hebbian covariance rule:

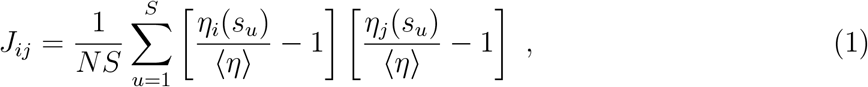

where *N* is the number of units with full recurrent connectivity, and the tunnel of length *L* is discretized into spatial bins of width *L*/*S* centered in *s*_*u*_ (*u* = 1, …, *S*). Here, ⟨*η*⟩ denotes the average activity over all cells and all positions. The distribution of activity {*η*_*i*_(*s*)}, the spatial code to be stored in memory, is chosen to replicate the statistics observed in bats flying in a long tunnel.

We focus on three sources of variability, as illustrated in Fig. 1. Each unit is taken to have, along the tunnel, a variable number *M* of place fields, given by a distribution of mean ⟨*M*⟩ (we use, as for the experimental data, an exponential fit, *P*(*M*) ∝ exp(−*M*/*ζ*), see App.A.1 for finer details). Each field *k* has a Gaussian shape of width *d*_*k*_ and peak rate *p*_*k*_; both parameters are drawn from *log normal* distributions, which fit quite well the experimental data. First, we assign *d*_*k*_ such that ln(*d*_*k*_) is normally distributed with mean *μ*_*d*_ and variance *σ*_*d*_; then we do the same for *p*_*k*_ but, to be really close to the data and in particular reproduce the observed correlation *ρ* ≃ 0.29 between *d*_*k*_ and *p*_*k*_, we take ln(*p*_*k*_) to be normally distributed with mean 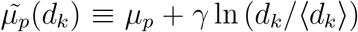 and variance *σ*_*p*_. This is a convenient formulation, as the overall mean remains the same, 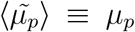, both distributions remain log normal, and by setting *γ* ≃ 0.5 we reproduce the observed correlation *ρ* ≃ 0.29. Besides this weak correlation, in the “realistic” model we assume a random placement of the fields and Gaussian fields truncated at *±d*/2, to accurately replicate neuronal recordings (see App.A.1 and Fig.2G). By using the specific parameters *ζ* = 4.7, exp *μ*_*d*_ = 4.8*m*, exp *μ*_*p*_ = 4.7*Hz, σ*_*d*_ = 0.575, *σ*_*p*_ = 0.884 and *γ* = 0.5, the field distributions obtained from the “realistic model” are nearly identical to the original recordings (see Fig. 1). We thought that inserting *γ* = 0.5 to reproduce the correlation *ρ* = 0.29 may be excess accuracy, and initially proceeded with a simplified analytical approach by setting *γ* = 0, but were proven wrong, as discussed below. Recently, moreover, Burak and colleagues have been able to show that precisely that degree of correlation would arise naturally in a model in which the place fields result from random Gaussian processes (Yoram Burak, personal communication).

**Figure 1.**
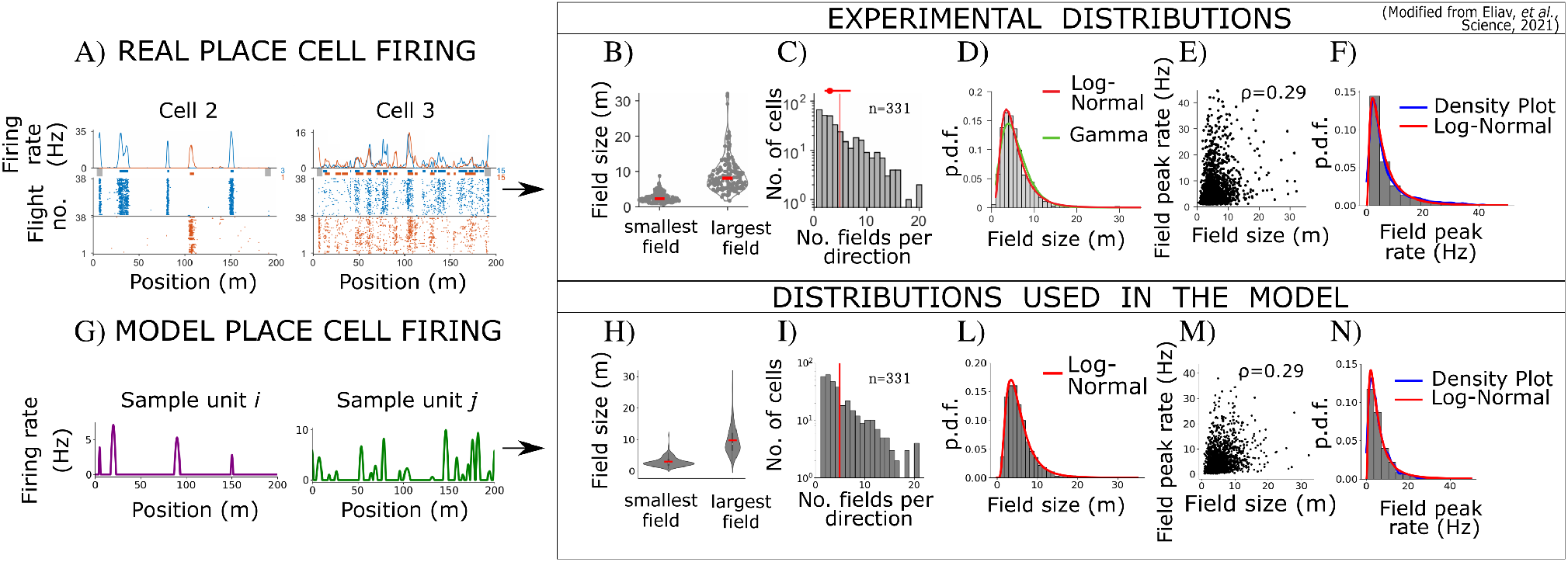
Our “realistic” model (second row) considers distributions similar to those observed in CA1 (first row). Panels A)-D) are reproduced with permission from [15] for the sake of comparison. Panel F) was obtained from the experimental data in E), kindly given us by the authors. A) Firing rate profiles of two neurons and G) of two sample units whose number of fields, peak firing rates and fields dimension where sampled from the model distributions. B)H) Distribution of the smallest and largest field sizes per neuron (B) or per model unit (H) (including those with at least 2 fields). C)I) Distribution of the number of place fields per neuron (C) or per model unit (I) in one direction. The bar at 20 includes all numbers above 20, and the average number of fields is ⟨*M*⟩ = 4.9 in both distributions. D) Distribution of fields sizes as obtained in experiments, the log normal parameters of the fit (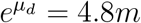, *σ*_*d*_ = 0.575) coincide with those used in the model (L). E)M) Scatter plot of field size versus peak firing rate for each field, as obtained in experiments (E) or with our algorithm (M). Note the non-zero *ρ* = 0.29 Spearmann correlation coefficient between the two measures. F) Distribution of the experimental peak firing rates, fitted with a log normal distribution parametrized by (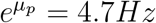, *σ*_*p*_ = 0.884) N) Distribution of peak firing rates as obtained with the algorithm, using the same parameters. The sample distributions in H)-N) were obtained with *N* = 331 units.

Once it has been released from external inputs and connections are consolidated, the network evolves under its recurrent connectivity only. We call {*V*_*i*_(*t*)} the activity of each cell *i* at time *t*, during such recurrent dynamics. The discrete time evolution equation for neural activity reads

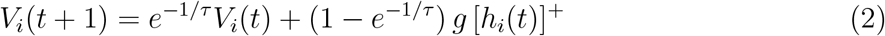

where [*h*]^+^ = max(0, *h*), *τ* < 1 is the time scale for converting the rapid spiking of real neurons into gradually changing firing rate dynamics (in units of simulation time steps), *g* is a fixed gain parameter, and

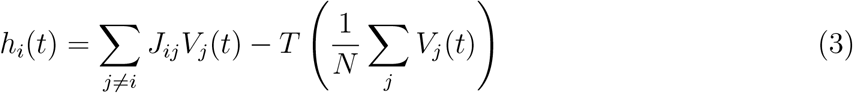

is the input current to unit *i* at time *t*, relative to a threshold *T* incorporating fast activity-dependent inhibition. We consider *T* (*v*) = 4*k*(*v* − *v*_0_)^3^ where *v*_0_ is the target mean activity. Note that these dynamics can be associated with an energy (Liapunov) function as described in App.A.2.

### 2. Dynamical encoding of position by neural activity shows three different regimes

To assess to what extent a spatial code has been established in the network memory, we estimate the cosine similarity between the configuration of activity 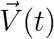 and the population vectors 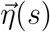 at all positions along the tunnel

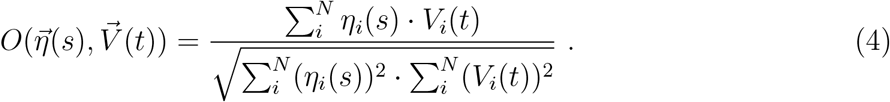

This overlap, bounded from above by unity, is a measure of the quality of position encoding. The set of overlaps 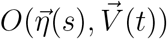 as *s* spans the environment defines the overlap profile at a given time *t*. If it is a bump-like profile peaked in *s*^*^, then the activity robustly codes for this position. Conversely, when the profile is not spatially tuned, the configuration of activity lies out of the attractor. By tracking this profile over time, we therefore obtain a characterization of the high-dimensional neural dynamics.

Initializing the network activity at some location *s*, i.e. 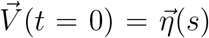 (but alternative assignments of the initial activity vector 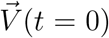 lead to the same results, see App. A.4 Fig. 9), we monitor the time course of the overlap profile. Three scenarios are observed:

- Continuous quasi-attractor (CQA): the bump is present at all times, and slides along the manifold until a fixed point is reached (Fig. 2A). The bump remains tightly packed throughout the motion, see Fig. 2D.
- Fragmented Manifold (FM): the bump deforms, gradually spreads out (see Fig. 2B and App.A.3 for details) and then re-localizes, as if teleported elsewhere, until it reaches a fixed point (Fig. 2E). Informally speaking, at some point the manifold has effectively ceased to exist, or to be relevant for population dynamics.
- Non localized (NL): the bump rapidly grows in width and ceases to be localized (Fig. 2C) until a non-informative fixed point is reached, spread over the whole environment (Fig. 2F).

By spanning a wide range of values of the field-defining quenched parameters *ζ, σ*_*p*_, *σ*_*d*_ we are able to obtain a phase diagram, locating the regions associated to the CQA, FM and NL regimes, shown in Fig.2H-M as three (*σ*_*p*_, *σ*_*d*_) planes for three different *ζ* values. To draw these planes we employ three types of measurements. First, from the dynamical runs, we extract the proportion of vanished manifold, as the fraction of initial positions from which population activity jumps somewhere else, see Fig. 2 (H, I, J) and Fig. 3 (C, G). Second, at the fixed points, we compute the alignment of the direction of instability of the fixed points with the direction of the presumed 1D manifold, as illustrated in Fig. 3 (B, F); as well as, third measure, the spread of their overlap, as depicted in Fig. 2 (K, L, M). Further details can be found in App. A.3.

**Figure 2.**
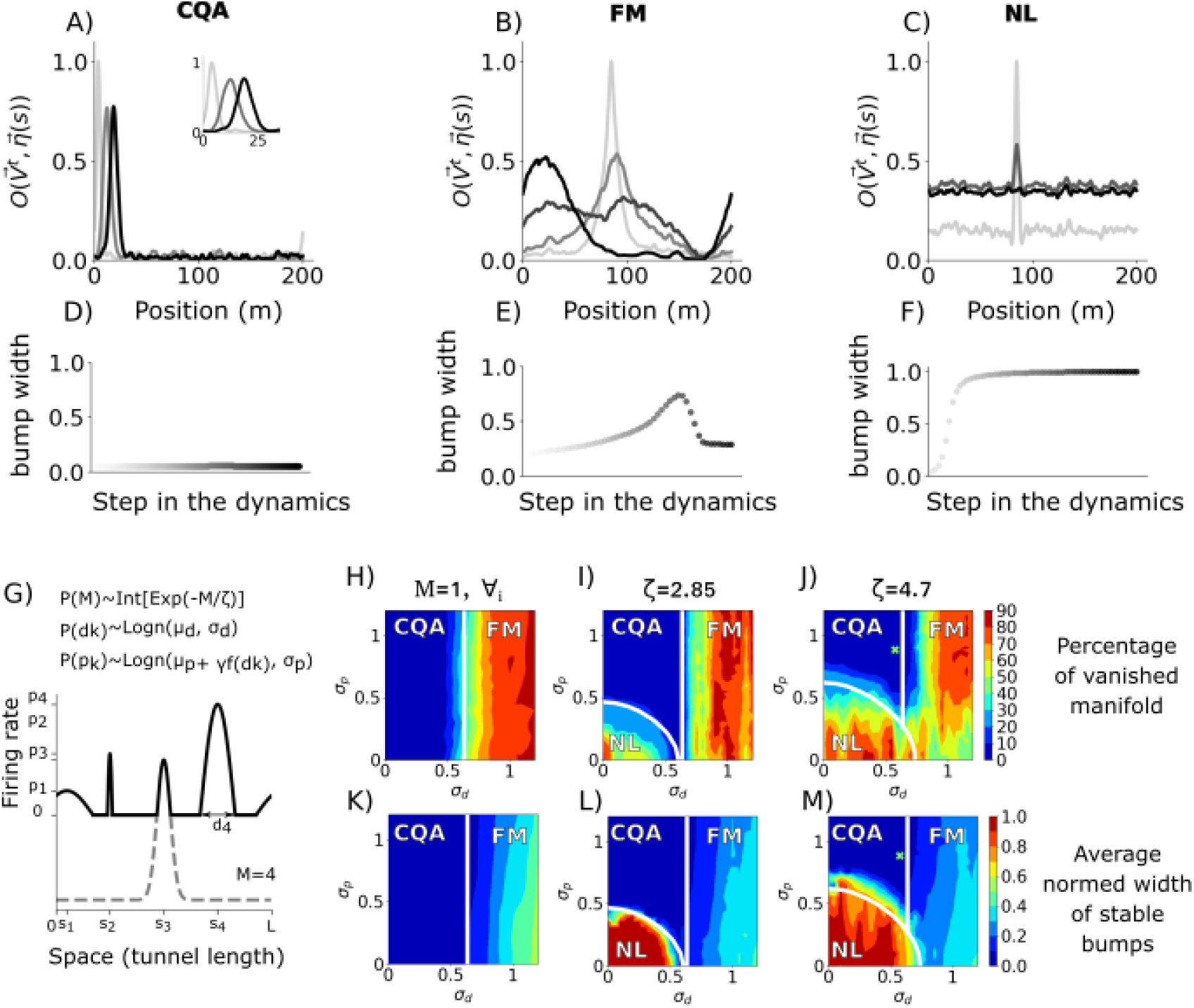
In the “realistic” model, a Continuous Quasi-Attractor can be established only within an intermediate variability range. A-F) Numerical characterization of the three regimes. A-C) Sample dynamics, as they reach the fixed points starting from a configuration of activity localized on the manifold, illustrated by the overlaps *O*(*η, V* (*t*)) with the patterns to be stored. The selected times *t* are indicated by colors ranging from light gray (initial condition) to black (fixed point). The dynamical states in the CQA regime slide on the manifold (A) – the inset shows a zoom-in; the ones in the FM regime B) jump outside and re-enter it elsewhere; while the ones in the NL regime C) get delocalized. D-F) Estimation of the standard deviation around the center of mass of the overlap profiles (after removing ripples below *O*(*η, V*) = 0.1). G) Black line: firing rate map of one sample unit with *M* = 4 fields. Each field *k* is characterized by a field peak rate *p*_*k*_ and diameter *d*_*k*_ drawn from the log normal distributions where f(*d*_*k*_) is defined in App.A.1, while the position of the center *s*_*k*_ is drawn from a uniform distribution. Dashed gray line: as an example, below the third field we show the profile of the full untruncated Gaussian used to generate each field *k*. H-M) Phase diagrams depicting, in the *σ*_*p*_-*σ*_*d*_ plane, both numerical and analytical results for increasing average number of fields (*M* ≡ 1 for H)K); *ζ* = 2.85, ⟨*M*⟩ ≈ 3.4 for I)L); *ζ* = 4.7, ⟨*M*⟩ ≈ 4.9 for J)M)). H)I)J) show the average fraction of the quasi-attractive manifold which vanishes, K)L)M) show the average normalized activity bump width of the fixed points (see Supplementary Material for details about these and other measures, like the average number of fixed points and their average sparsity). Green crosses indicate the parameter values from the experimental results. White curves are derived analytically (see text). Variability parameters: A)D) *σ*_*d*_ = 0.4, *σ*_*p*_ = 0.4, *M* ≡ 1, *N* = 8000; B)E) *σ*_*d*_ = 0.9, *σ*_*p*_ = 0.9, *M* ≡ 1, *N* = 8000; C)F) *σ*_*d*_ = 0.2, *σ*_*p*_ = 0.2, *ζ* = 4.7, *N* = 4000. *H* − *M* obtained by interpolation (with 26*x*26 data points, only 12*x*12 for H,K). A data point is averaged over 2 − 5 different quenched realizations of the network, each probed with 50 simulations of the dynamics, *L*/*S* = 0.1,*N* = 16000 *g* = 17 *k*_*b*_ = 300. Different reasonable values for these parameters do not affect much the boundaries between the regimes, while larger *N* values make them sharper (see also Fig.3 and App. A.4).

**Figure 3.**
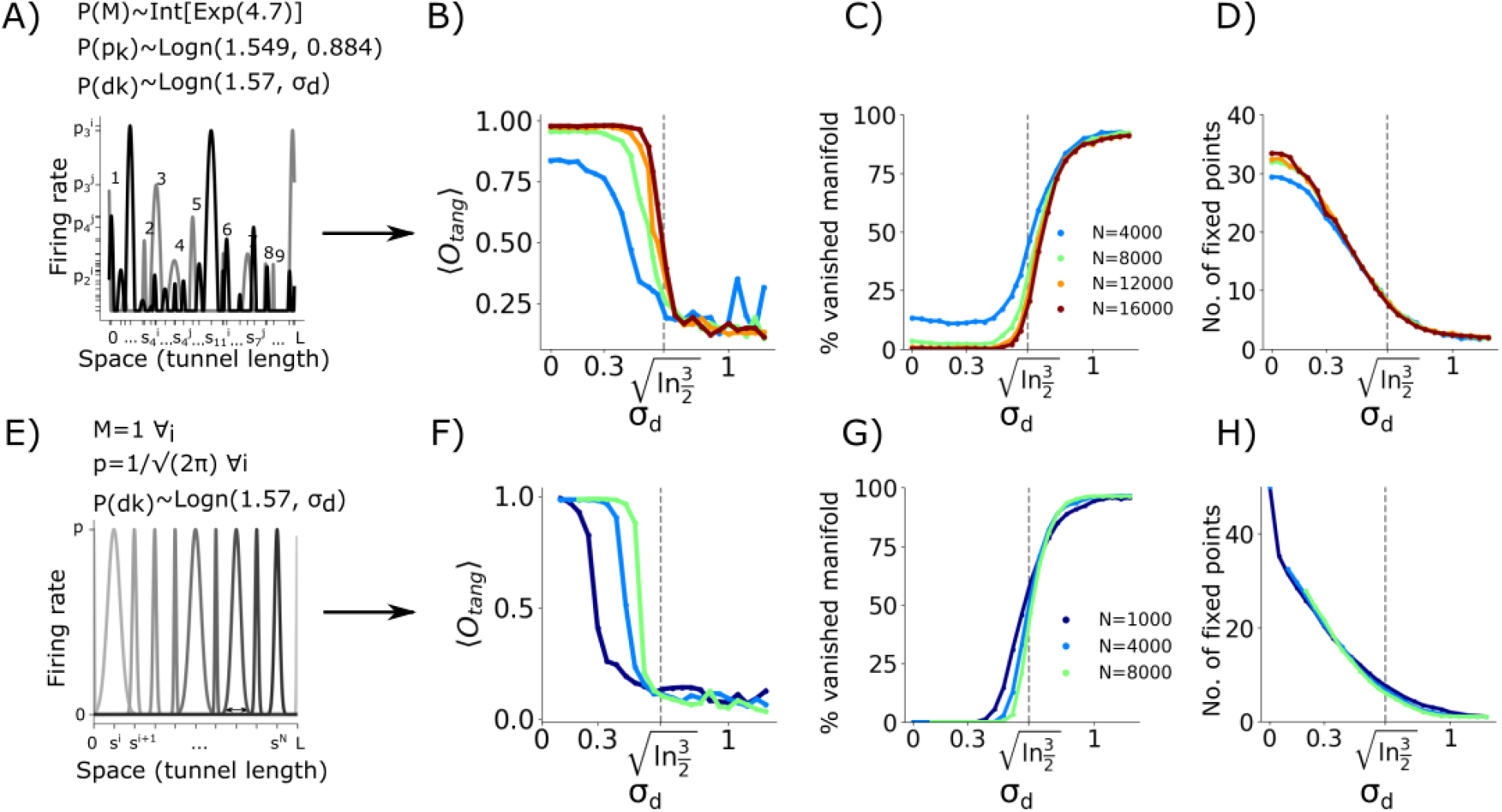
The teleportation boundary can be studied with a much reduced model. The upper row shows results obtained using the realistic model, with *σ*_*p*_ and *ζ* set as in the experimental distribution (*ζ* = 4.7, *μ*_*p*_ = 1.549, *σ*_*p*_ = 0.884). The lower row shows those obtained with the simplified model (*M* = 1, all peak rates set at 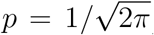). A) Multiple irregular fields of two sample units (gray/black) with *σ*_*d*_ = 0.5, numbers indicate the field label of one unit. E) Single fields of 10 sample units in the reduced model with *σ*_*d*_ = 0.87. B)F) ⟨*O*_*tang*_⟩ quantifies in the range 0 − 1 the alignment of the most unstable eigenvectors with the direction of the manifold (see App.A.3 for details); The curves indicate the 0.25 quantile (75% of the overall data lie above the line) for the realistic (B) and the simplified (F) models. C)G) Fraction of runs in the two models in which the overlap profile deforms, indicating a jump outside the quasi-attractive manifold, taken as a proxy for its break-up (see App.A.3); D)H) In contrast, the average number of fixed points does not depend on the size of the system and decreases smoothly with *σ*_*d*_. The vertical lines at 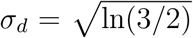 show the analytical estimates derived from the simplified model. A-D) simulations are run following the algorithms described in the Appendices and all parameters except *σ*_*d*_ are set to model experimental results. Each point on each curve is obtained averaging over 25-90 quenched realizations of the network (fewer when the system size is larger), and 50 simulations per network (20 for the second row), each initialized at a different {*η*_*i*_(*s*)}, with *s* regularly spaced every 4*m* along *L*.

**Figure 4.**
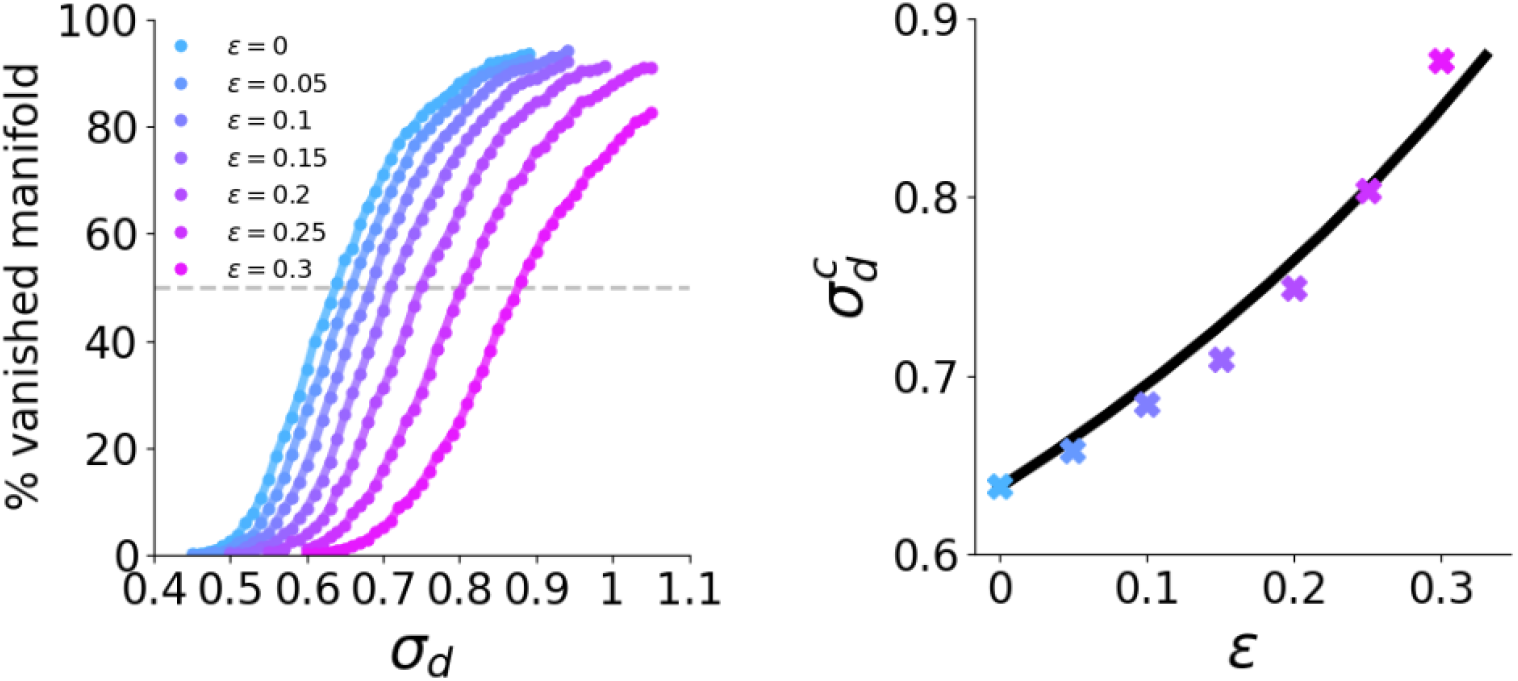
The *ϵ*−expansion yields critical values 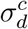 consistent with those estimated from simulating the reduced model. Left) As in Fig. 3 C-G, for increasing *ϵ* values. Right) crosses: 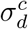 estimated at 50% of the curves on the left. The black curve is obtained analytically, as described in the text; Parameters: individual uniformly distributed purely gaussian fields, *N* = 4000, *g* = 20, *k*_*b*_ = 300, *L*/*S* = 0.2.

**Figure 5.**
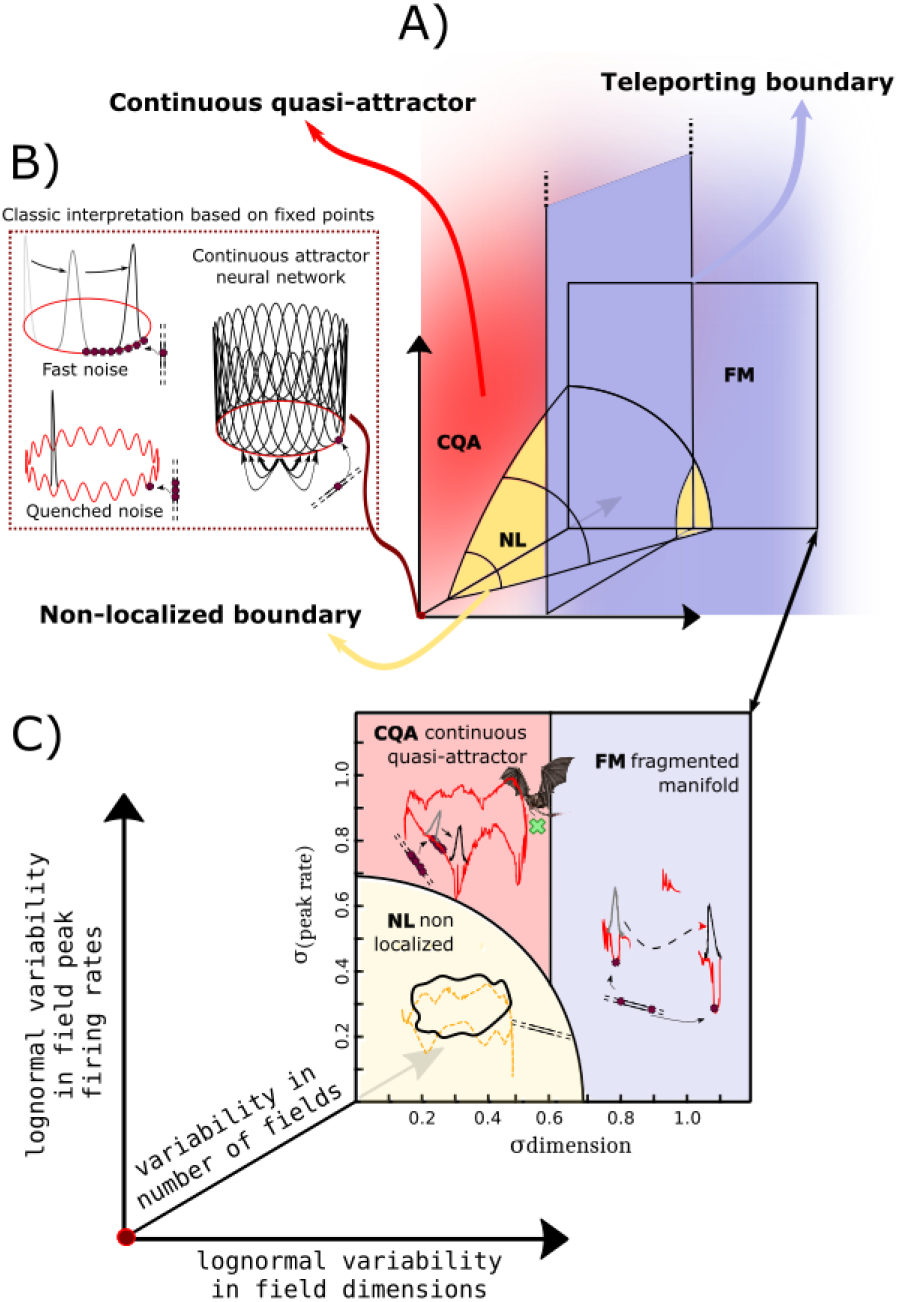
Interpretation of the phase diagrams: a continuous quasi-attractor model for memory storage and retrieval. A) Rough sketch of the three dimensional phase diagram, with axes labels as in C), schematizing the results reported in Fig.2. Three regimes are separated by two boundaries. In the continuous quasi-attractor regime (red), the manifold of solutions, representing different locations in space, is attractive to all directions but one: an external cue can drive the network states towards the manifold, and spatial memory is expressed as distinct bumps of activity sliding along the non-attractive direction. In the teleportation or Fragmented Manifold regime (purple), the manifold effectively vanishes: external cues can only drive the dynamics towards a few residual fixed states, unable to represent space, as dynamics do not smoothly flow along the manifold. In the Non-Localized phase (yellow), the fixed points are not localized: hence no cue can retrieve spatial memory. B) Sketch of the idealized continuous attractor (right) which can only emerge, in the continuous quasi-attractive phase, at the origin, i.e. in the unrealistic condition of single, equal and regularly positioned fields. The parallel lines on the bottom right of the cANN symbolize a 1D track: each fixed point in the continuous attractor neural network (purple cross) represents a memory of one position in the environment (matching purple cross in the track). The presence of mild fast/quenched noise (its effects on the cANN are schematized on the two left sketches) downgrades the precision of spatial memory. C) Sketch of the three regimes in a plane corresponding to the distribution in the number of fields observed in the recording in bats ([15]). Memory retrieval is preserved as long as the dynamical flow is aligned with the manifold. This occurs in the red region, which includes also the firing rate and field width statistics observed in the recordings (green cross). Memory capability suddenly deteriorates beyond each boundary (determining the inability to retrieve anything, or possibly only a few locations). In C) and B) the closed curves (red/yellow) are intended to sketch the energy of the quasi-attractive continuous manifold. Gray-black bumps indicate the overlap profile which one can calculate with the {*η*(*s*)} at each step of the dynamics. The yellow dashed manifold, instead, represent a putative manifold which may exist but is not reached by the dynamics.

**Figure 6.**
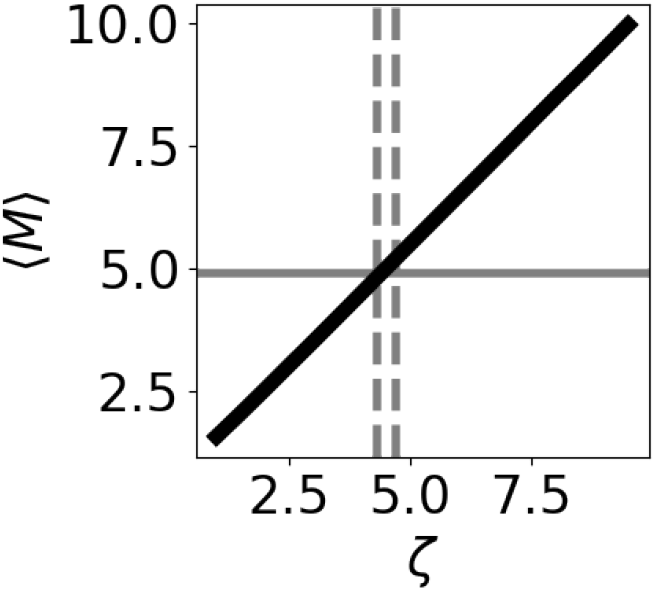
Numerical estimation of ⟨*M*⟩ drawing random numbers from the exponential distribution Prob(*M*; *ζ*) (see text) under the described constraints. The horizontal black line shows the experimental average value ⟨*M*⟩, while the vertical dashed lines show a range of arbitrary choices of *ζ* (4.3¡*ζ*¡4.7) to obtain, roughly, a matching average ⟨*M*⟩=4.9 as in the experiments.

**Figure 7.**
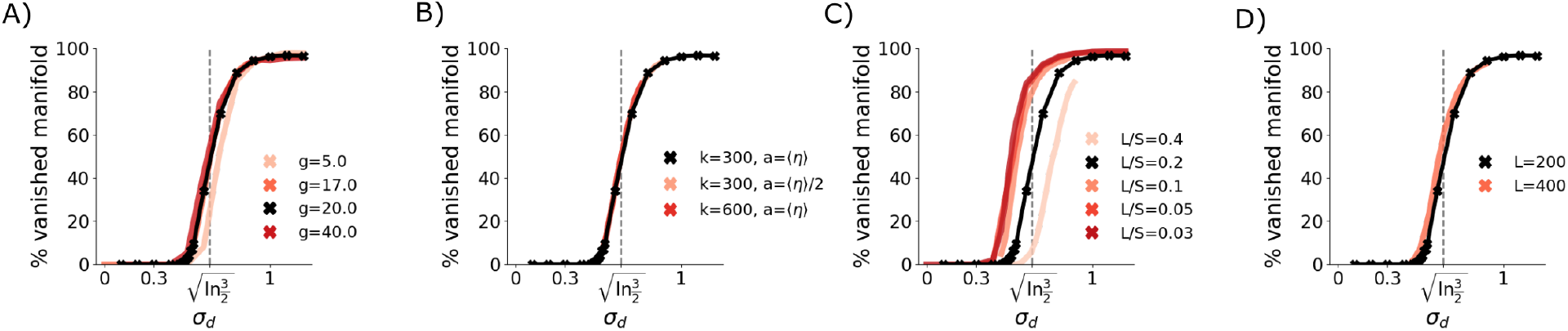
We have tested several parameters around a prototypical set (black) – space length *L* = 200 discretized in *S* = 1000 bins, threshold constant *k* = 300, limit average activity *a* = ⟨*η*⟩, gain *g* = 20. We see that none of these parameters affects substantially the critical level of variability in field size, *σ*_*d*_, at which the manifold appears to break up. Although the optimal granularity *L*/*S* would be ∞ 0.1 in our simulations, we keep *L*/*S* = 0.2 for computational reasons, to allow for smaller and faster simulations.

**Figure 8.**
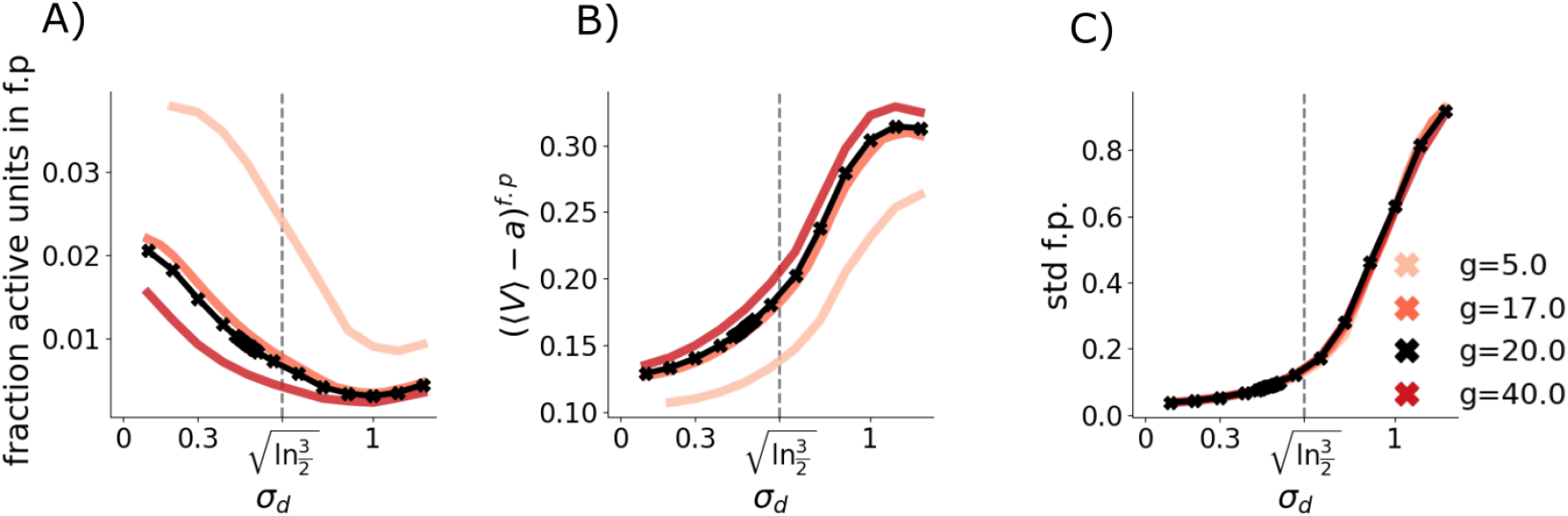
The parameter which mostly affects the fraction of active units in the fixed points (A) and the capability of the network to set the average activity (B) is the gain, without affecting the size of the retrieved bump (C).

**Figure 9.**
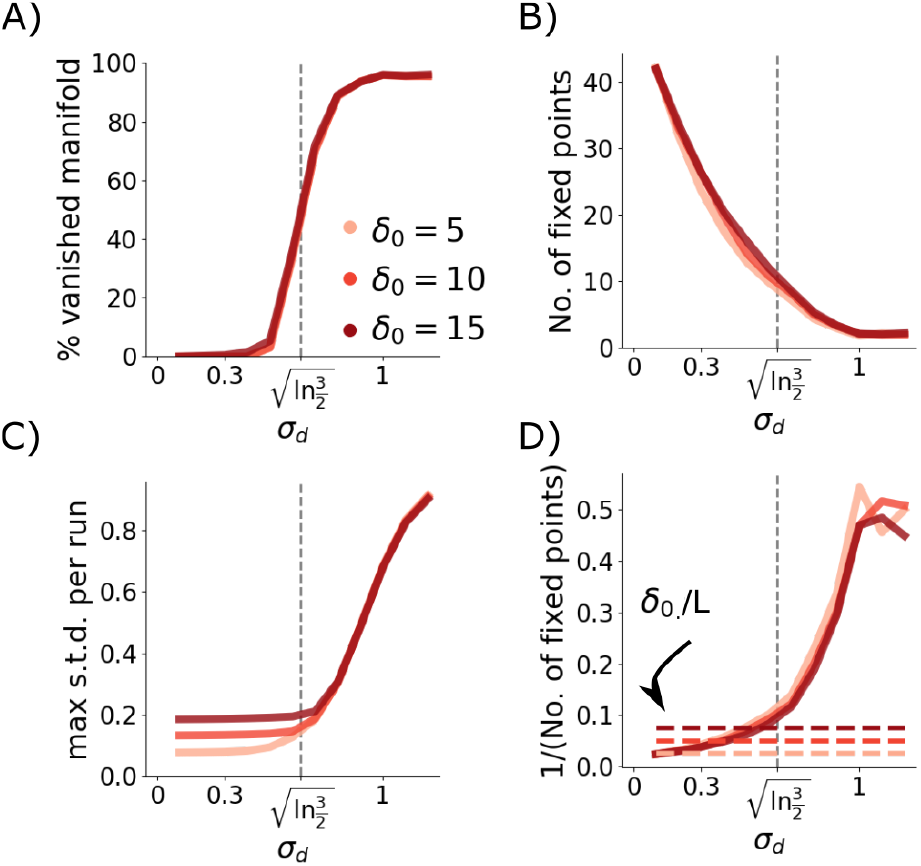
In the simplified model, i.e. when all units have single fields, uniformly distributed along the environment (see Fig. 3E), we can verify that the width of the initial condition does not affect much whether activity jumps outside the manifold. We take the initial activity of each unit *i* whose field is centered in *s*_*i*_ to be a gaussian function of the distance from a bump center *s*^*^: 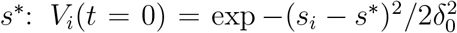, with a spread parametrized by *δ*_0_. Having tested for several network realizations and center positions, we find that the 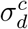 value at which the manifold appears to vanish (A) as well as the number of fixed points (B) do not depend on the width of the initial activity bump. The maximal width of the bump during the dynamics (C) does depend on the initial width *δ*_0_, but only up to the critical value 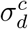, at which value the overlap profile becomes, at some point in time, very large, as it comes to reflect the jump outside the manifold. Even when the average distance between the fixed points is larger than the width of the initial condition (D) – dashed lines represent the standard deviation in unit values, i.e. some units at the fixed point were inactive in the initial bump – the convergence towards the fixed points is smooth, below 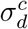, while it wanders around in activity space outside the manifold, above 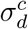.

Some general features characterize this summary of the simulation results, as seen in Fig. 2 and in Fig. 3:

- The CQA regime is found in a central region of the (*σ*_*d*_, *σ*_*p*_) plane, when *ζ* > 1.
- For *σ*_*d*_ above a critical value that appears to be almost independent of *σ*_*p*_ and *ζ*, the network abruptly transitions to the FM regime, becoming effectively unable to represent, based on memory, almost any position along the tunnel (Fig. 2 H, I, J). This critical value is above, but close to the experimental value from CA1 recordings, *σ*_*d*_ = 0.575. The exact location of the boundary between the CQA and FM regimes is also not sensitive to the precise values of the gain and threshold, which only affect the sparsity and the fraction of active units at the fixed points (see App. A.4). Note the contrast between the sharp increase with *σ*_*d*_ in the proportion of starting positions from which activity is eventually teleported, see Fig. 3 C (and App. A.4), and the smooth decrease in the number of fixed points, which in itself would not suggest that the network is in a distinct phase.
- When units have multiple fields, and both *σ*_*d*_ and *σ*_*p*_ are small, approximately within a circular boundary (which depends on *ζ*), population dynamics always delocalizes, and the network can be said to have entered, again abruptly, the NL regime.

### 2. Boundaries of the phase space region where continuous quasi-attractors emerge

Obtaining analytical boundaries would enable us to rigorously prove that the continuous quasi-attractors break up through a real phase transition rather than a mere cross-over to a different regime, and also to predict where the boundary would be for other values of the various parameters. Yet, it is challenging, due to the non-stationarity of the phenomenon. Nevertheless, we have derived satisfactory analytical estimates of the boundary positions (see Fig.2, H)-M), white curves) using signal-to-noise analyses, as reported below.

#### Quasi-circular region of non-localization

When units have multiple fields, the fact that with limited variability in their width and peak rate network activity gets “smeared” over the entire length of the tunnel, as shown in Fig.2F, indicates that the competition among the fields fails to produce a winner. Intriguingly, adding variability in the distribution of peaks and diameters promotes order, i.e. it allows the network to better differentiate among the different fields. To quantitate the competition, consider a simplified model with *N* units that have exactly *M* fields each (if *M* varies, like in the simulations of the realistic model, in which it is a random variable taken from a quasi-exponential distribution parametrized by *ζ*, we can later consider an average across *M* values). Only the strongest field 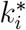 of each unit *i*, if one at all, should participate in localizing activity along the tunnel, with the other fields *k* = 1, …, *k*^*^ − 1, *k*^*^ + 1, …, *M* contributing just noise. Since the competition is within the spatial preferences of individual units, we can focus on one, drop the index *i*, and take *k*^*^ = 1 without loss of generality. If all the fields have the same shape, the “’strength” of each field field *k* is proportional to the product of its width and peak rate

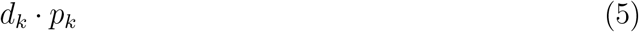

and the average signal and noise contributed by a generic active unit are given by

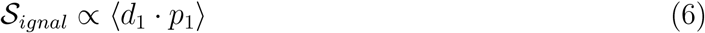

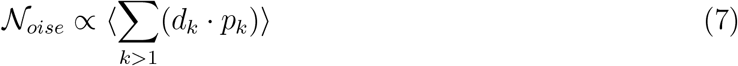

In the model, *d*_*k*_ and *p*_*k*_ are taken each from a log normal distribution, with a correlation between them parametrized by *γ*, as in Eq. (21), so if we assume that active units reflect those distributions, we can change variables

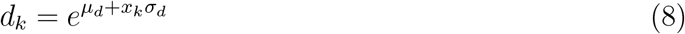

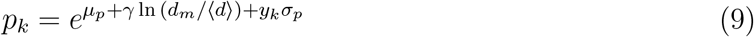

and perform the integrals as shown in App.A.5, simply by restricting the domains of integration to enforce the constraint that *k*^*^ = 1 be the strongest field. The result is

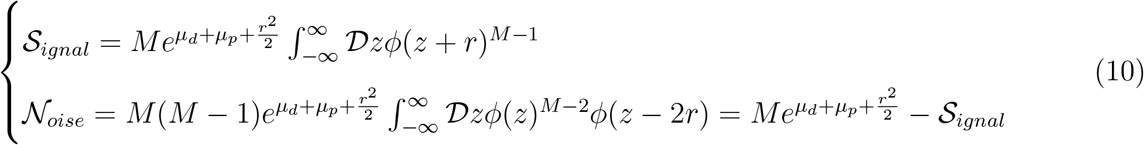

where

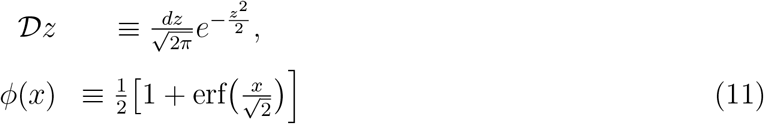

is the cumulative Gaussian integral and

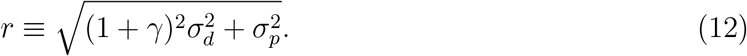

If now active units have each a different number *M* of fields, one can evaluate numerically, for a given *ζ*, the weighted averages ⟨𝒮_*ignal*_(*M*)⟩_*M*_ and ⟨𝒩_*oise*_(*M*)⟩_*M*_ .

We have empirically found, as reported in App.A.5, that we can approximately describe the quasi-circular boundary by setting the signal-to-noise ratio equal to the ratio between the mean peak rate and its standard deviation, scaled down by a factor *A*. In a formula, the boundary, which depends on *ζ* (or, in general, on ⟨*M*⟩), would comprise the values 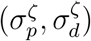 satisfying the equation

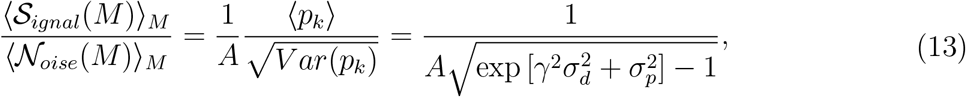

which is derived and written in full in App.A.5.

It is not obvious that the equation above roughly describes a quarter of a circle in the (*σ*_*p*_, *σ*_*d*_) plane. In fact, we had initially set the r.h.s simply to 1, in which case the l.h.s, which depends only on *ζ* and *r* (see App.A.5), would be equal to 1 exactly on a quarter circle, *provided γ* = 0. A circular boundary is *not*, however, what one finds simulating the simplified model with *γ* = 0, i.e., uncorrelated distributions for *d*_*k*_ and *p*_*k*_, as shown in App.A.5 Fig.10. The boundary is circular, or approximately so, only when one uses for the simulations correlated distributions with *γ* close to 0.5, which leads to a Spearman correlation near the experimentally observed value *ρ* ≈ 0.29 [15].

**Figure 10.**
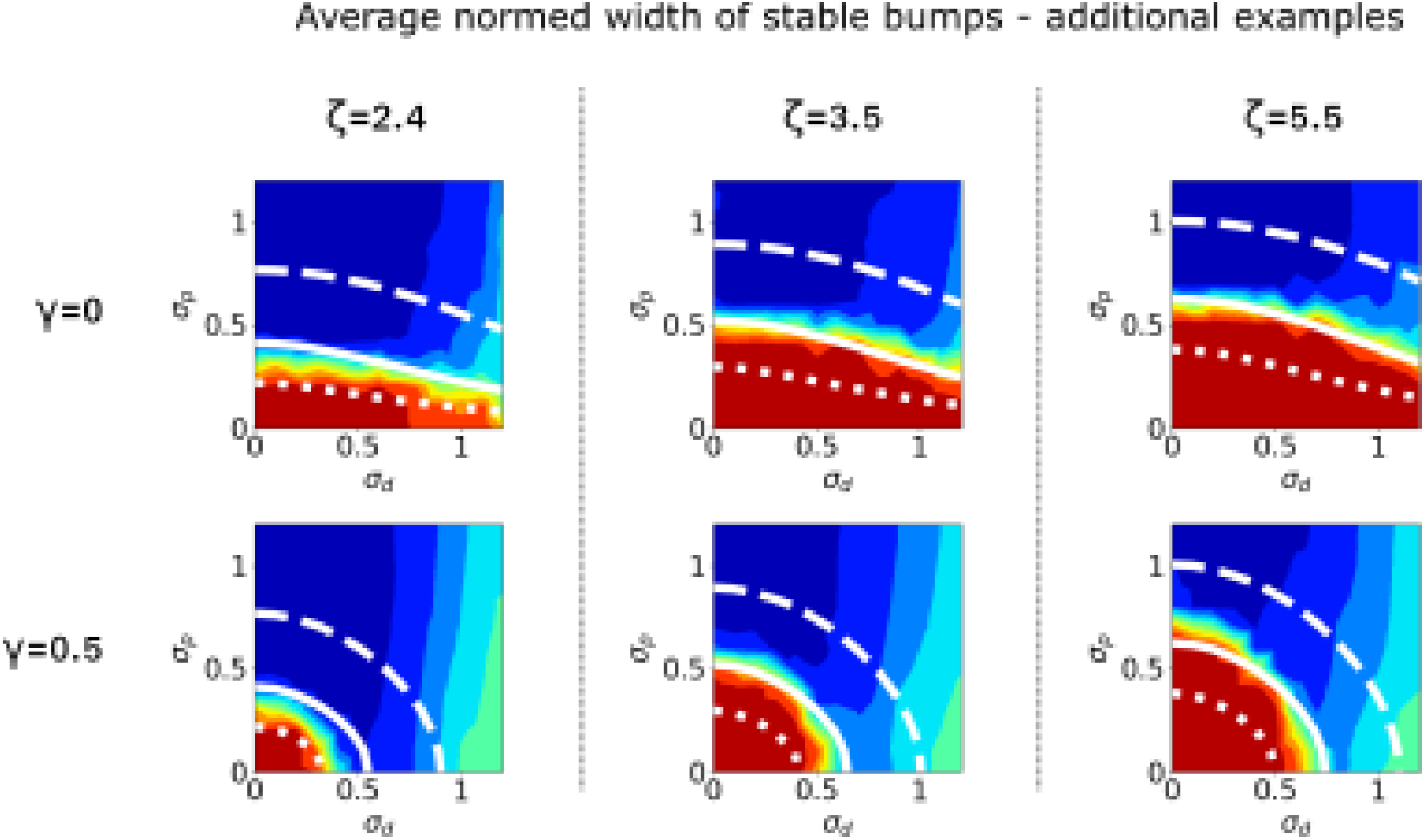
Complementary to Fig.2, we plot the average width of the fixed points for three different values of *ζ*, setting the average numbers of fields, with the respective analytical estimates of the boundary between the non localized regime and the localized one (white curves) for three different values of *A* = 1 (dashed), *A* = 3 (solid), *A* = 7 (dotted). In the first row we show the results one would obtain in the absence of correlation between peaks and diameters, *γ* = 0, while in the second one with the correlation determined by *γ* = 0.5, leading approximately to a Spearman correlation *ρ* = 0.29, as in the experimental results Eli+21.

**Figure 11.**
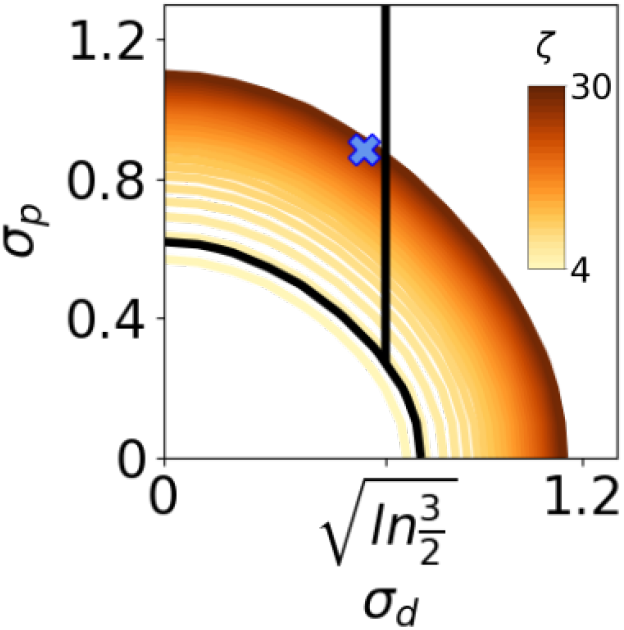
With increasing *ζ*, the boundary of the non-localized regime expands while remaining quasi-circular, until at *ζ* ≃ 27 it reaches the values *σ*_*p*_ and *σ*_*d*_ measured in the bat experiments [15]. For *ζ* slightly larger, and the experimental value for *σ*_*p*_, there is no longer a CQA range of *σ*_*d*_ values (see text).

This led us to write the r.h.s. as in Eq.(13) above. Whichever value one chooses for the factor *A* (*A* = 3 in Fig.2; see App.A.5 for further details), it can be verified numerically that this form of the r.h.s. restores a quasi-circular boundary. Its effective radius increases with *ζ*; and therefore the region inside the circle, where the noise exceeds the signal, also expands, and population activity increasingly does not localize, when the mean number of fields per unit increases – a trend which has interesting implications (see below).

#### Vertical boundary towards fragmented manifolds

We note that, in the simulations, the vertical boundary appears to be independent of *ζ* and nearly independent of *σ*_*p*_. We introduce then another simplified model, *ad hoc* to study this other boundary. In this new model, each unit is characterized by a *single* field with a fixed peak rate. These fields exhibit purely Gaussian shapes and are uniformly distributed across the environment. The only source of variability among the units lies in the widths of their fields, which follow a log normal distribution as in the more realistic model (Fig. 3 A and E). Simulations of this reduced model reveal that, similar to the realistic model, there is a critical value 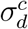 at which the fraction of vanished manifold abruptly increases and the alignment of the direction of instability of the fixed points with the manifold abruptly decreases (Fig. 3). Prior to this critical value, the qualitative behavior during the dynamics remains the same as that described for the realistic model.

The activity jumps because, for large 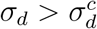, the disparity in the width of the fields makes it likely that a few of the widest fields will happen to be centered far away from the initial position *s*. If they still get activated and start suppressing the narrower units centered around *s*, the self-reinforcing process can lead to a jump. Critical to the self-reinforcing nature of the process is that the largest fields exert a disproportionate influence with respect to the narrow ones; this is because in the Hebbian plasticity rule used in the “realistic” model (Eq. 1) the normalization of both pre-and postsynaptic factors is by the *average* activity level across units, ⟨*η*⟩.

To treat this simplified model analytically, we can consider the average (square) fluctuation in the inputs coming from units at a location *s* far away from the current activity bump, due to the chance concentration at that location of units with large fields. This is now the (square) noise, due to other units and not, as for the quasi-circular region, to the other fields of the same units, and we propose to estimate it as

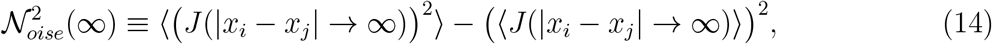

and it can be compared with the signal, estimated as the average input from the units at that location,

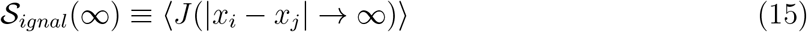

where we have considered that the calculation, as reported in App. A.6, simplifies greatly if the signal 𝒮_*ignal*_(*s*) and the noise 𝒩_*oise*_(*s*) are calculated at *s* → ∞, after taking also the limit *L* → ∞. With the Hebbian rule in Eq. (1), 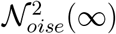 depends on *σ*_*d*_ and one finds

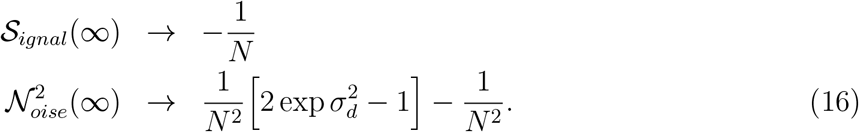

𝒮_*ignal*_(∞) is negative, of course, but the fluctuations quantified by 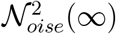 can overcome it, if *σ*_*p*_ is large enough. Setting 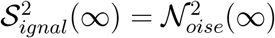, one obtains 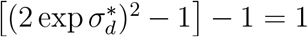, which has the solution

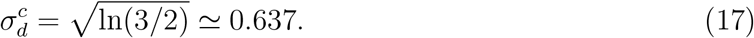

Numerically, this solution is indeed close to the critical value estimated from the simulations (white vertical lines in Fig. 2, dashed vertical lines in Fig.3).

The validity of the analysis above is supported by its ability to locate the transition point for a class of plasticity rules, in which the couplings are given by

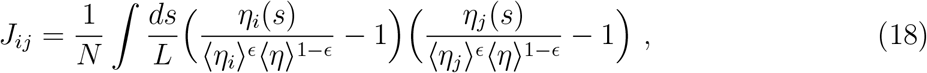

where 0 ≤ *ϵ* ≤ 1. For *ϵ* = 0, the original Hebbian rule in Eq.(1) is recovered. For a small and strictly positive *ϵ*, the normalization of the coupling *J*_*ij*_ now slightly depends on the neurons *i* and *j* through their average activities ⟨*η*_*i*_⟩ and ⟨*η*_*j*_⟩. With this plasticity rule, applicable only in the reduced model, the units with wider fields (which can now be defined, as each unit has only one field in this model) would exert an influence over a wider range of locations, but a bit weaker at its peak than the units with the narrower fields.

The calculation of the critical 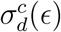 which yields 𝒮_*ignal*_(∞) = 𝒩_*oise*_(∞) (see App. A.6 for details) gives, for small *ϵ*, values close to those obtained in simulations using the unusual *ϵ*-normalization in the plasticity rule of the reduced model, as shown in Fig.4. We observe that, as *ϵ* grows, the transition arises for more inhomogeneous place field sizes. This result is expected based on the mitigating role of the neuron-dependent normalization of the coupligns in Eq.(18). In the limiting case of *ϵ* = 1, one does not in fact observe any jump, or teleportation, when simulating the network defined by the reduced model, indirectly confirming the self-reinforcing mechanism described above. Notice that this limiting case corresponds to an implausible plasticity rule: the normalization specific to the activity of each unit would vary if, for example, the environment is enlarged a bit, and some units acquire new fields, leading to substantial changes in the associated couplings.

## 3. Discussion

While the storage of multiple regular place maps – as distinct continuous attractors in the same connectivity matrix – induces quenched disorder, fragmenting each attractor but not destroying it completely, up to the capacity limit Bat+98,pap+07,Cer+13,Mon+13,Mon+14, Mon+15; we have seen that the storage of a single irregular map can already be enough to dispose of the continuous attractor as such, also in its weaker quasi-attractive version, in which the continuity is in the flow rather than in the fixed points. We have mapped through simulations a three dimensional phase diagram, spanning the variability in the number, dimension and peak firing of the fields. We find that in a portion of such a diagram, including the level of variability observed in CA1 recordings in bats, a continuous quasi-attractor can be established by model Hebbian plasticity; or rather it *emerges*, if regarding Hebbian plasticity as un unsupervised, self-organizing process. Memory is functional in this region, and place cell maps can effectively implement a quasi-attractive continuous manifold. Note that in the network model memory is functional but already fragmented, in the CQA regime, so that when external stimulation is absent, such as during sleep or rest periods, one would expect also the real network to be able to replay only fragments, one after the other. Thus, only fragments of the tunnel near the fixed points, which are more stably stored in memory, might be reached during spontaneous, rambling dynamics. This phenomenon has in fact been reported in the analysis of CA1 resting state recordings [54].

We see that the three-dimensional functional portion of the phase diagram has two distinct boundaries. If the variability of the diameters exceeds a critical value 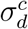, the quasi-attractive manifold breaks up, only a few fixed points are retained, and they attract activity initially localized all over the environment. This boundary is independent of *ζ*. On the other hand if, given *ζ* (and the correlation parameter *γ*), the joint variability in field width and peak rate is below a quasi-circular boundary (described approximately by a critical value 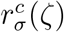), population activity delocalizes and its attractor dynamics cannot express spatial memory.

The surprising finding is that the regime boundaries seen by simulating the finite size and quasi-realistic model, which includes several rather arbitrary parameters, are matched quite closely by those estimated analytically, with rather tentative procedures, in the “thermodynamic limit”. The latter end up being parameter-free pure numbers; one dependent only on *ζ*, i.e. roughly on the mean number ⟨*M*⟩ of fields per unit (see App.A.1), the other not even on that, Eq.(17). Note that *ζ* is expected to scale linearly with the length of the tunnel, or in general the size of the environment, whether in CA1 or in CA3. As a result, for any given level of peak rate variability *σ*_*p*_ (and correlation parameter *γ*) there will a maximum size of the environment which can be stored in memory, when the two boundaries collide and the “acceptable” *σ*_*d*_ range shrinks to zero. It is given implicitly by the relation

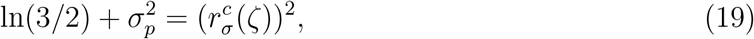

which with the parameters from the CA1 recordings in [15] would yield ⟨*M*⟩ ≃ 30, that is, *L* ≃ 1.3*km*, a surprising low limit for the length of a memorable tunnel.

The average over directions and the question of whether it is legitimate to neglect place field directionality, in our model, highlight the fact that the regime boundaries obtained do not depend on the dimensionality of physical space. Although derived after considering the statistics of place cell activity in a uni-dimensional tunnel, they should hold for spatial memory in 2D and 3D, as well, if the same log normal statistics applies. In that case, the limit given by Eq.(19), once the proper *σ*_*p*_ value is inserted, would amount to an absolute storage capacity limit for spatial memory, given as a concrete value for the size of the environment, unlike the rather abstract limits expressed as a maximum number of charts [11, 12].

Perhaps the incumbent capacity limit may explain why the real CA1 network appears to sit so close to a disabling phase transition (after taking into account all the caveats mentioned above). Alternatively, one might speculate that being on the verge of breaking its attractors is advantageous for a hippocampal network in other ways, e.g. to facilitate the acquisition of new maps or yet-to-be-explored portions of a familiar tunnel, or to reap the benefits sometimes ascribed to self-organized criticality [55].

## Acknowlegments

We are grateful to Nachum Ulanovsky, Tamir Eliav and their colleagues for extensively sharing and discussing their findings, as well as to other participants in the “M-Gate” collaboration. Kwang Il Ryom helped with numerics, and recently Yoram Burak clarified a point about their model of random spatial codes.

## Author contribution

FS and AT conceived the study, and FS designed, ran and refined multiple layers of computer simulations. FS, RM and AT made repeated efforts at interpreting the results and describing them analytically, and all co-authors jointly wrote the paper.

## A Appendices

### A.1 Algorithm for data generation

We take as inspiration the experimental results recently published by [15], and focus on three main sources of variability: the number of fields, the field size and the field peak rate.

#### A.1.1 Number of fields

[15] have observed that place cells recorded in large environments can have up to 20 fields (Fig. 1C) spanning from small to large ones (Fig. 1B,D) with an average of 4.9 fields per cell in each flying direction.

In order to simulate the distribution shown in Fig. 1C, we consider that the number of fields *M* per each unit is randomly drawn from an exponential distribution with probability function Prob(*M*; *ζ*) ∝ exp(−*M*/*ζ*) under the constraint that zero values, or values above *M* = 21, are not accepted. This constraints leads to a distribution of higher average than *ζ*, and we find that setting *ζ* = 4.7 results in ⟨*M*⟩ ≈ 4.9. In Fig.6 we report the numerical relation between the parameter *ζ* of the probability Prob(*M*; *ζ*) and the effective average number of fields ⟨*M*⟩.

#### A.1.2 Field shape and size

We consider each field *κ* to be characterized by: i) the position *s*_*κ*_ of its center, ii) its linear dimension, or effective diameter, *d*_*κ*_ and ii) its peak firing rate *p*_*κ*_. Please note that we assume periodic boundary conditions, such that fields are effectively lying on a ring of dimension *L* = 200*m*, which we depict, through most of the article, as open rather than closed, for ease of visualization. Further, we assume that each field can be modeled as a Gaussian bump, centered at *s*_*κ*_, with dimension *d*_*κ*_ = 2*σ* and maximal height *p*_*κ*_ (Fig.2G).

The activity of a unit *i* at a certain position *s* is thus given by:

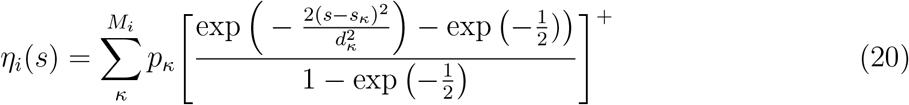

where *M*_*i*_ is the number of fields assigned to unit *i* and [ ]^+^ sets all negative values to zero.

For the sake of reproducing the statistics measured in experiments we do not allow fields to overlap, and we constrain the sum of the dimensions of all fields belonging to a unit to be less than (*L* − 3*M*_*i*_)*m*, where *L* = 200*m* is the size of the environment and the last subtraction facilitates finding appropriate random positions for the fields on the ring.

We randomly draw the size *d*_*κ*_ of each field from a log normal distribution ℒ(*μ*_*d*_, *σ*_*d*_) with exp *μ*_*d*_ = 4.8*m* and *σ*_*d*_ = 0.575, as resulting from the fit to the experimental data reported by [15] (Fig. 1D).

#### A.1.3 Peak rates

The peak firing rate distribution, in the experimental recordings, can be roughly fit with a log normal distribution ℒ(*μ*_*p*_, *σ*_*p*_) with parameters exp *μ*_*p*_ = 4.7*Hz* and *σ*_*p*_ = 0.884, which we estimated from the data obtained by [15] (reproduced in Fig. 1E) kindly given us by the authors.

In order to introduce the correlation seen in the experimental recordings (Fig. 1E), given a field with a certain dimension *d*_*κ*_ we define the specific mean

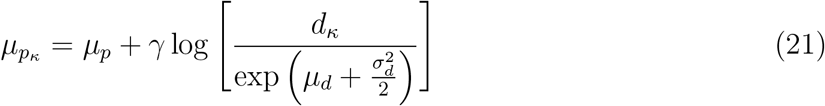

with *γ* = 0.5, and draw the first guess 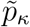 of the peak firing rate corresponding to that field from a log normal distribution with that mean. Then, we prevent having fields with peak firing rate much higher than 40Hz by using instead of 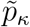 directly, its non-linear transform *p*_*κ*_, inspired by those used in phonology to transform Hz into Bark. In particular we obtain a modified peak firing rate of a field *κ* as:

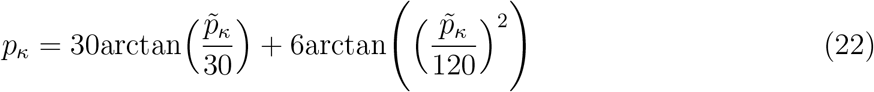

These functions, overall, lead to distributions satisfactorily agreeing with the recordings, as shown in Fig. 1 E-F and M-N.

Remarkably, as reported in the main text, the main results do not appear to depend on the details of the distributions and are general also in simpler systems, without the constraints (*M* < 21, modified *p*_*k*_, Gaussian fields truncated at *±d*_*k*_/2). The latter, which specify the “realistic model”, are solely required to reproduce satisfactorily the observations in [15].

### A.2 Energy and Hessian functions

Given the system described in Eqs. (1), (2), (3), we define a quantity

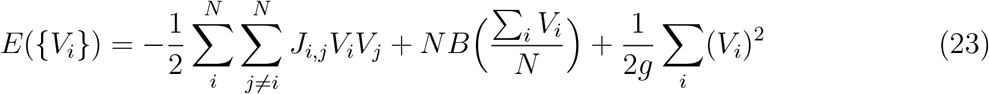

with *dB*(*v*)/*dv* ≡ *T* (*v*), i.e *B*(*v*) = *k*(*v* − *v*_0_)^4^. For any unit *i* above threshold, *V*_*i*_(*t*) > 0, we have

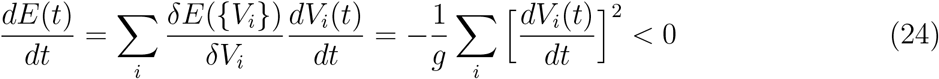

showing that *E* behaves as an energy function in the hyperquadrant *V*_*i*_(*t*) > 0, as introduced in [56]. Then the Hessian is

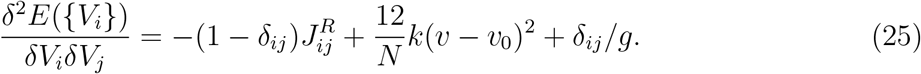

In Eq. (25) 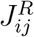 is the weight matrix *J*_*ij*_ restricted to the units above threshold: columns and rows of those below threshold are removed.

### A.3 Measures of loss in attractivity

One can think of different approaches to quantify the loss of attraction, in the remaining *N* − 1 directions, of the quasi-attractive continuous manifold. Here, we use three measurements.

1. **Proportion of dynamical runs which jump**: Given a large number of simulations starting at different 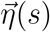, with *s* spanning the whole length *L*, we take the proportion which exit the manifold, i.e. those in which the localized bump does not slide continuously, as a measurement of the percentage of the manifold which has vanished. In particular, given a system we run a number ≥ 50 of dynamics each starting from a certain configuration 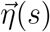, such that the overall length of the manifold is uniformly sampled. 50 dynamics are enough, for the variable systems we are studying, to reach each fixed point configuration, typically several times. At each dynamical step, we estimate the position of the center of the bump *s*^*a*^. If at step *t*+1 the position of the center *s*^*d*^ has moved farther than a certain, arbitrarily set, distance (i.e |*s*^*a*^−*s*^*d*^| > 20*cm* – or less), we check that the overlap values progressively decrease from *s*^*d*^ to *s*^*a*^, i.e. that 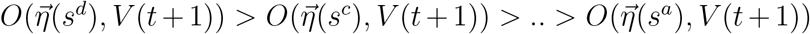. If this does not occur for more than a given small number of discrete positions (corresponding to < 1*m* in the model) we regard these dynamics to have jumped outside the manifold. If instead it occurs for less than 1*m* we consider this effect as a “physiological” ripple of the bump.
2. ⟨***O***_**tang**_⟩: Given the complete set of residual fixed points on the manifold we evaluate, for each, the eigenvector corresponding to the smallest eigenvalue of the Hessian matrix (Eq. (25)) and estimate its cosine similarity with the direction of the manifold. This quantity, which we name *O*_tang_, equals 1 only if the eigenvector closest to instability is aligned with the manifold. In particular, in order to calculate the cosine similarity between the eigenvector closest to instability and the direction of the manifold, one needs to estimate the direction of the manifold around each fixed point, 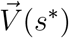, where *s*^*^ is, as elsewhere, the position at which the profile of the overlap between the memories and the activity state is maximal. This is not trivial, as we have precise access only to the fixed point configurations and to estimate the direction numerically one needs to know the activity coding for the position just preceding the one at the fixed point and take the difference. To do so, once we have estimated *s*^*^ we run the dynamics with a long time scale *τ* (see Eq.(2)) starting from an initial condition 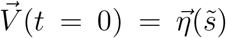 where 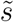 is closer to *s*^*^ than to any other fixed point position. Then we estimate the direction as the difference between 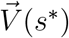 and the configuration of activity, incurred along the way from 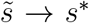, having maximal overlap with the closest discretized position *s*. One can take either the average among all dynamical configurations of activity having maximal overlap with the position before *s*^*^, or the one occurring just before the maximal overlap slides towards *s*^*^, and results are very similar.
3. **Bump width:** Given a configuration of activity, whether this is a dynamical one or a fixed point, we can estimate its localization on the manifold as the normalized standard deviation of the center of mass of its overlap profile with {*η*(*s*)}. The closer this value, once normalized, is to 1 the more the configuration of activity is spread on the manifold and the less it is localized. To make this quantity informative we remove sources of noise in the overlap profile by thresholding it to an arbitrarily set value of 0.1. Fig.2D-E-F shows the estimated bump width during the dynamical evolution reported in Fig.2A-B-C.

In particular, given a vector 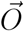, where each entry is the overlap *O*_*s*_ = *O*(*η*(*s*), *V*) (Eq. (4)) with the respective quenched pattern, the center of mass is given by

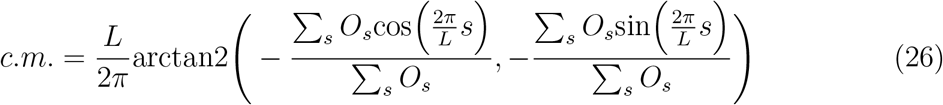

where *L* is the total length and the trigonometry is used to implement periodic boundary conditions. The standard deviation around the center of mass, instead, can be calculated by first estimating the vector 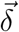 of the minimal distances between each *s* and the center of mass (keeping in mind periodic conditions) and then as

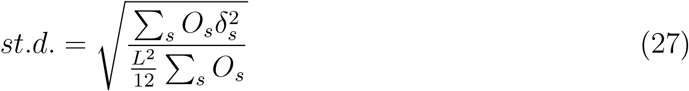

As the irregularities of the shape of the bump outside the localized region substantially increase the standard deviation, thus reducing the information regarding how localized is the bump, we remove all irregularities below 0.1 (we simply subtract 0.1 by all overlap values and send to zero those which become negative).

### A.4 Independence of the vertical boundary from the model parameters

The quantities we define to estimate the breaking of the continuous quasi-attractor do not depend much on the details of the model, as suggested by Fig.7.

### A.5 Analytical calculation estimating the quasi-circular boundary

Consider first the case that all *N* units have exactly *M* fields each. If *M* varies, like in the simulations, in which it is a random variable taken from an exponential distribution with mean *ζ*, one can later take suitable averages. Assume that only the strongest field 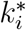 of each unit *i* participates in the localized bump, and the other fields *k* = 1, …, *k*^*^ − 1, *k*^*^ + 1, …, *M* contribute as noise. Let us drop the *i* symbol and focus only on one unit.

We define the strength of each field *k* as

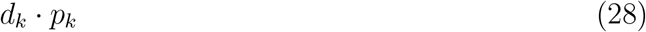

where *d*_*k*_ is its diameter and *p*_*k*_ its peak firing rate.

With these assumptions, the signal and the noise can be estimated as

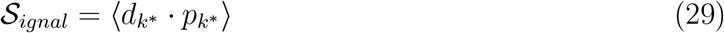

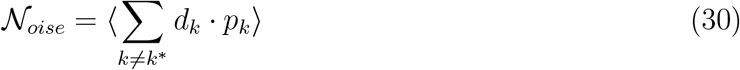

We take *d*_*k*_ and *p*_*k*_ to be independently drawn from two log-normal distributions, the latter depending on the specific value taken by *d*_*k*_ in the former. We express this through a change of variables

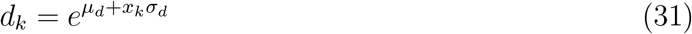

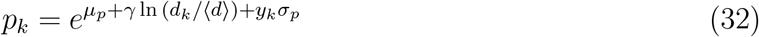

such that the probability distribution of each realization *x*_*k*_, *y*_*k*_ is normally distributed, and given by

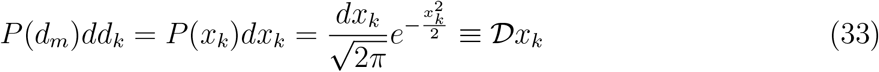

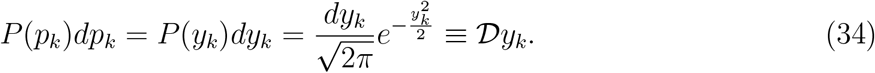

One can calculate the average signal and noise by considering that the strongest field is the first, i.e. *k*^*^ = 1, then

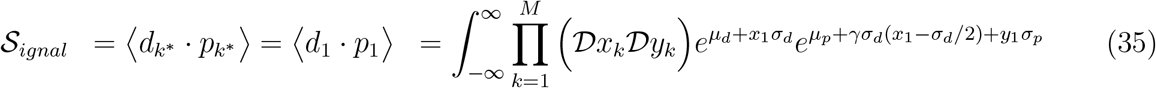

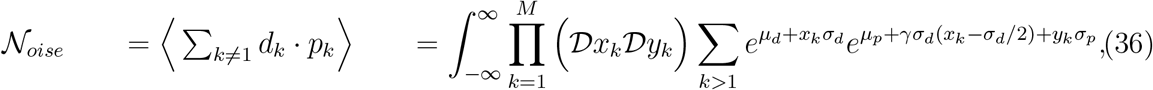

where in order to properly set the integration interval one should hold on to the integral for *k* = 1 while carrying out first the ones for *k* = 2, .., *M*.

Let us also consider that, in the expression for 𝒩_*oise*_, all terms *k* = 2, …, *M* have equal distributions, so one can consider *k* = 2 as the prototypical one and multiply by (*M* − 1) the result:

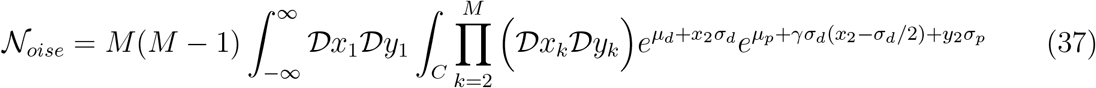

nd similarly for the signal

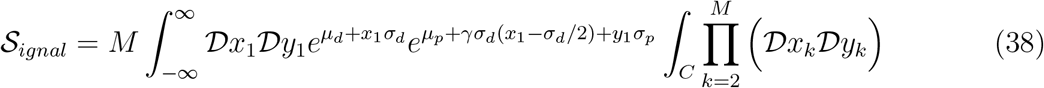

where *C* determines the combination of values (*x*_*k*_, *y*_*k*_) satisfying the condition that all fields *k* are less strong than *k*^*^ = 1, i.e., for ∀*k* ≥ 2

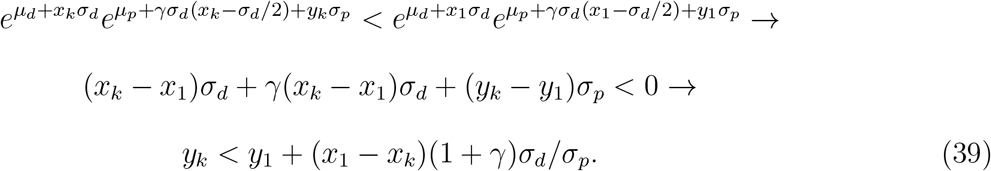

Note that the sum 𝒮_*ignal*_ + 𝒩_*oise*_ yields an even simpler expression, as it is not subject to the same condition; a fact we use below as a check.

One has two types of internal integrals (for *k* ≥ 2) in the expressions above. Both can be handled by rotating the variables to

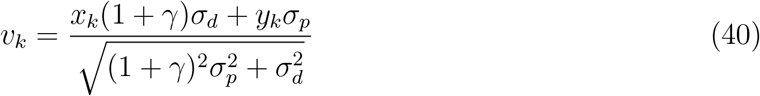

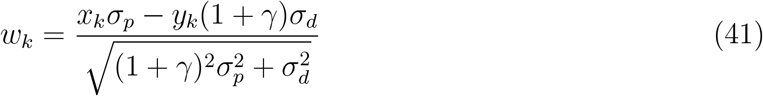

(note that the Jacobian remains |*J*| = 1), so that the condition on the integration domain reduces to *v*_*k*_ < *v*_1_, while *w*_*k*_ can be integrated from −∞ to +∞. One has

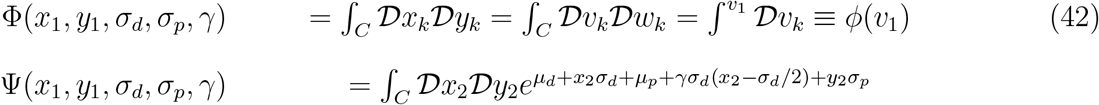

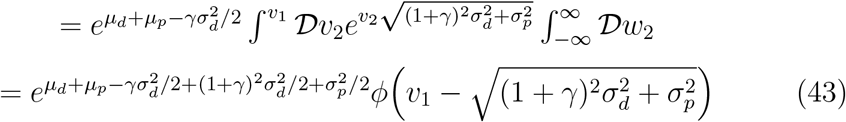

where

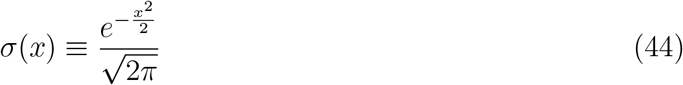

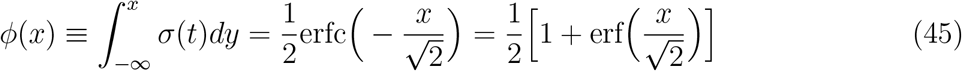

and to complete the last passage one has changed variables again to 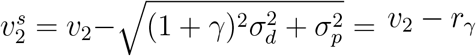, having defined the effective, *γ*-dependent radius

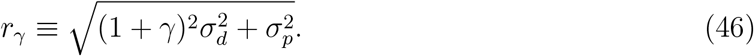

Then one can write

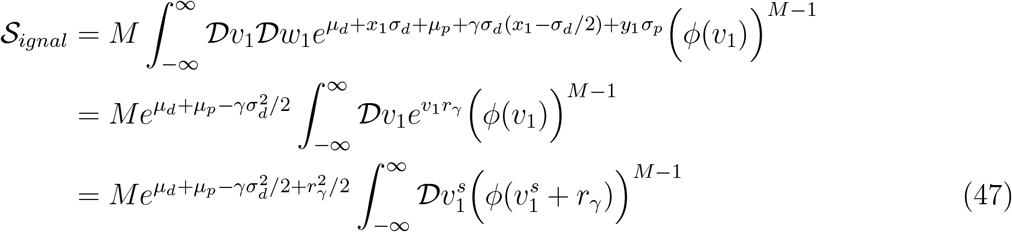

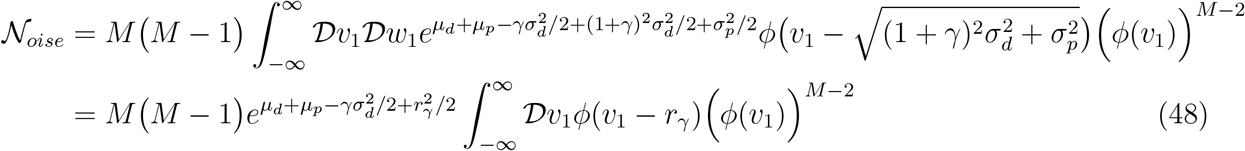

which are our final expressions for the signal and the noise. Note that integrating by parts the noise one has

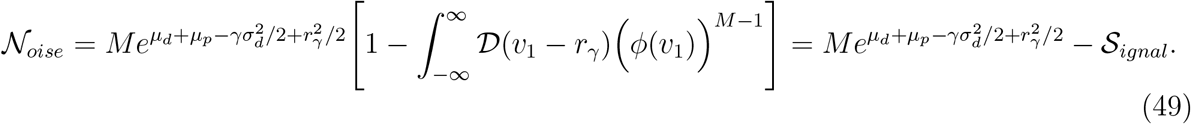

In conclusion, the signal to noise ratio, for fixed *M*, can be written as

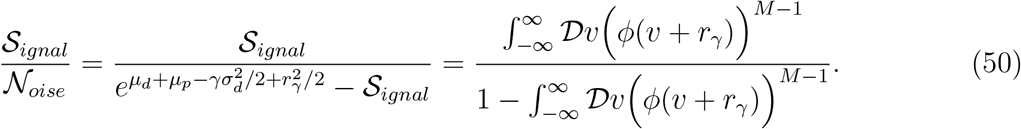

In the simulations, however, we have a variable *M*. We then define ⟨𝒮_*ignal*_(*M, γ, σ*_*d*_, *σ*_*p*_)⟩_*M*_ and ⟨𝒩_*oise*_(*M, γ, σ*_*d*_, *σ*_*p*_)⟩_*M*_ as the occurrence-weighted average of the signal and of the noise.

For *γ* = 0, both signal and noise depend on *σ*_*p*_ and *σ*_*d*_ solely through 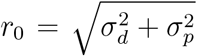, so obviously their ratio, set to any given value, describes a circle around the origin in our (*σ*_*p*_, *σ*_*d*_) phase plane. This is not, however the boundary between the localized and the non-localized regime observed in simulations with *γ* = 0, as shown in the App. Fig.10. The quasi-circular boundary is observed only when using, in the simulations, a value *γ* ≃ 0.5 reproducing the Spearman correlation *ρ* ≈ 0.29 observed experimentally by [15].

Thus the simulation results indicate that the average signal-to-average noise ratio should indeed be large, for activity to localize, but not relative to 1 or to a given set constant, rather to a function that brings the boundary back to being approximately circular, for *γ* ≃ 0.5 (and not for *γ* ≃ 0). We have found empirically, by trial and error, that the ratio of the signal and noise we have just estimated should be compared to another signal-to-noise ratio, that in the peak amplitudes, scaled down by a factor 1/*A*, where a suitable value is *A* ≈ 3. In a formula, the condition appears to be

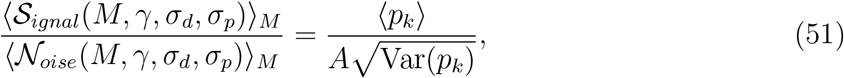

where the right-hand side does not depend on *ζ*, but it does depend on *γ* and *σ*_*d*_ implicitly, through the distribution of the peak values. We now proceed to calculate it explicitly. Given a correlation *γ* between field diameter and peak rate, we assume the diameters to be given by the standard log normal distribution, and the mean of the correlated log normal distribution for the peaks to be, for fields with diamak of the fields with diameter *d*

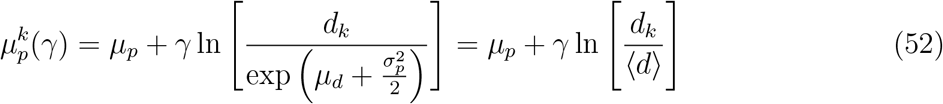

The average peak of the fields with diameter *d* is thus

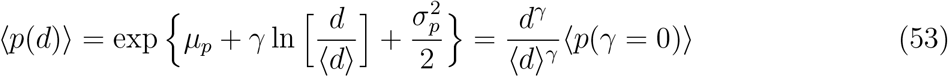

where 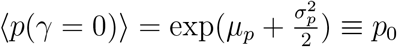.

The overall mean of the correlated peaks ⟨*p*_*k*_⟩, given the distribution of the diameters, is given by

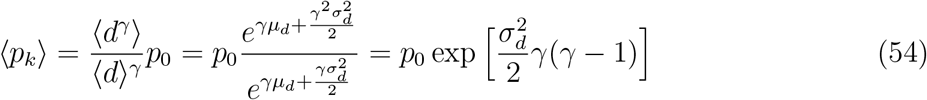

The variance of those field having diameter *d* is given by

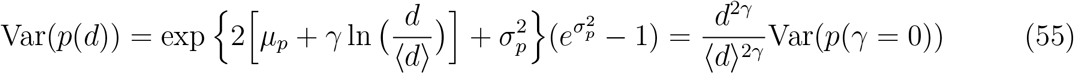

where 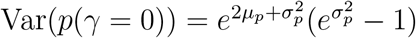; while the overall variance is given by

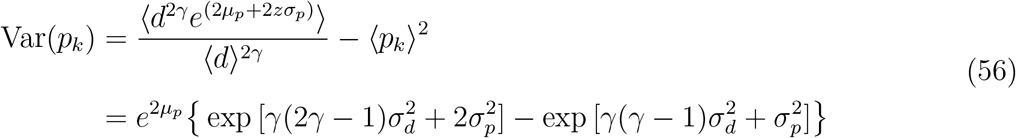

which under the square root and divided by ⟨*p*_*k*_⟩ gives

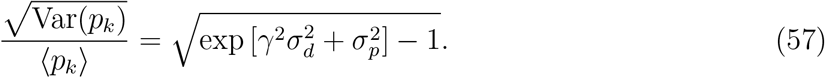

which leads to the expression in the main text Eq. (13).

### A.6 Analytical calculation estimating the vertical boundary

In the simplified model each unit has a single Gaussian field of fixed peak rate, with width *d* ≡ 2*r*

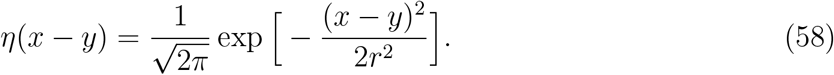

The diameters, or equivalently the radii, are lognormally distributed, so we can write

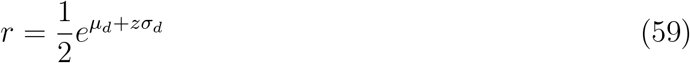

where *z* is a normally distributed variable of zero mean and unit variance.

We consider a slightly generalized form, with respect to Eq. (1), for the connection weight between a unit *i* with its sole field centered in *x*_*i*_, and radius *r*_*i*_ with a unit *j* centered in *x*_*j*_ with radius *r*_*j*_

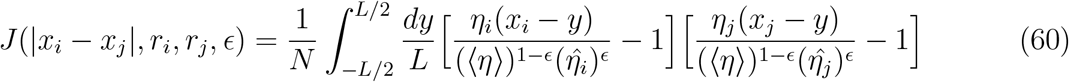

where the pre- and post-synaptic factors are normalized not simply by the overall mean of activation during learning ⟨*η*⟩, but rather by a geometric average of the overall mean with the mean specific for each particular unit, 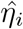. The exponent *ϵ* regulates the relative weight of the two, and for *ϵ* = 0 we have the standard case, which is the only logically consistent one in the full model in which each unit can have multiple fields. By considering, however, *ϵ* ≠ 0 and small, in the simplified model, we have a tool to validate the analytical prediction as to the position of the phase boundary, which turns out to depend on *ϵ*, and which can be checked against simulations carried out with *ϵ* ≠ 0.

The Gaussian field assumption implies that the “kernel” between any two units is also close to a Gaussian (minus a constant), of square width the sum of the two square widths

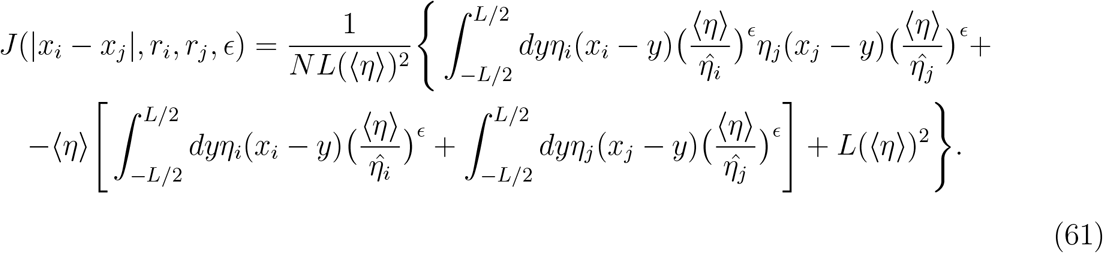

If we consider that the size of the ring *L* ≫ *r*_*i*_ for most units or effectively, we take the *L* → ∞ limit disregarding the lognormal tail of units with very large fields, we get

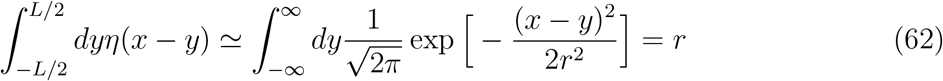

and

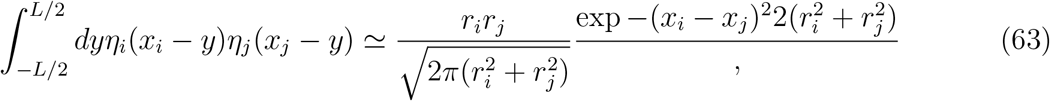

while obviously ⟨*η*⟩ ≃ ⟨*r*⟩/*L*.

Defining

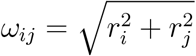

overall we have

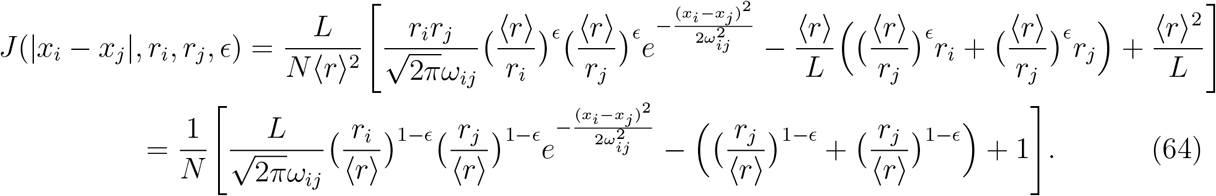

Consider now two units *i* and *j*. The connection weight between them fluctuates (around zero, the way we have defined it) due to the distance between their field centers *x*_*i*_ and *x*_*j*_ and also due to their radii *r*_*i*_, *r*_*j*_, as we see in Eq.(64). If the two field centers are very distant along the length *L*, however, the first term in Eq.(64) vanishes, and their interaction, which could potentially promote a jump of the network activity localized around *i*, to become localized around *j*, is on average negative. There are fluctuations, however, due to the values of the radii *r*_*i*_ and *r*_*j*_, and the jump may be promoted if the fluctuations overcome the negative mean. We can express the limiting condition as

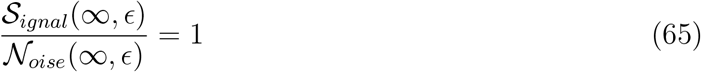

here we have defined the signal and noise at very large distances as

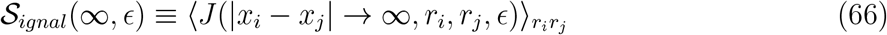

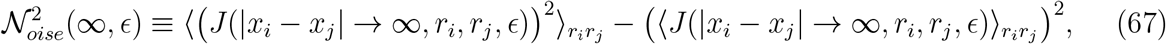

or, introducing the expression for *J*(|*x*_*i*_ − *x*_*j*_| → ∞, *r*_*i*_, *r*_*j*_, *ϵ*) and simplifying the notation

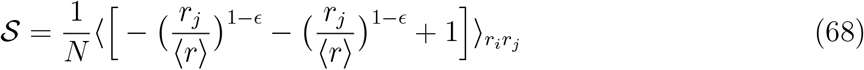

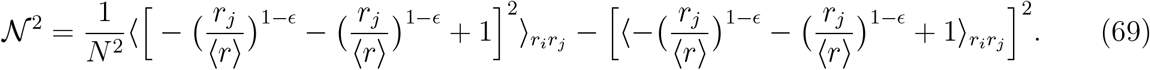

One is left, therefore with averaging powers of *r*_*i*_/⟨*r*⟩ whic, given their lognormal distribution, can be easily carried out giving

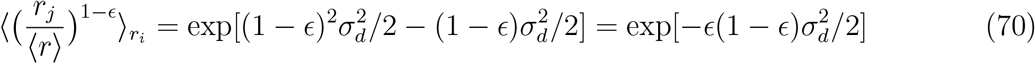

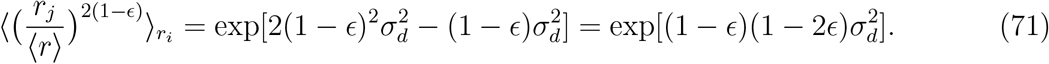

Collecting terms one finds

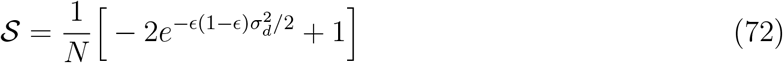

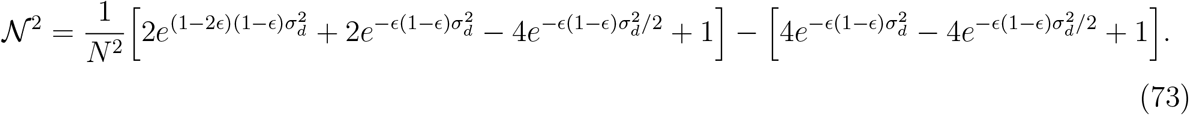

One can write the 𝒮/ 𝒩 = 1 limiting condition as 𝒩^2^ − 𝒮^2^ = 0, yielding

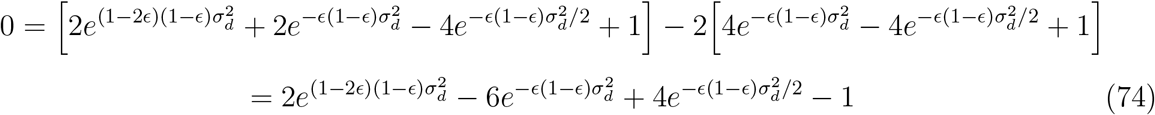

which in the case of practical interest *ϵ* = 0 gives

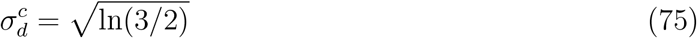

while for general *ϵ* gives the curve in Fig.4.

